# Optimisation of OptoDrum protocol for measuring optomotor response in juvenile & adult zebrafish

**DOI:** 10.64898/2026.05.20.720959

**Authors:** Rebecca Super, Bang V. Bui, Jiaheng Xie, Pamela Bou-Antoun, Leandro Scholz, Patricia R. Jusuf

## Abstract

Zebrafish (*Danio rerio*) are an important vertebrate model for vision and neuroscience research. In the larval stages, the aquatic species begins to elicit the optomotor response (OMR) to stabilize themselves in water — a behaviour that may be exploited in the laboratory to measure visual acuity. However, up to now, the measurement of the OMR in juvenile and adult zebrafish has been limited due to their behavioural complexity. Here, we optimize a protocol to assay zebrafish aged between 4 and 9 weeks-post-fertilization, by displaying sinusoidal gratings parallel to the zebrafish eye to elicit a robust OMR. We assessed the visual spatial-frequency tuning function of an environmentally induced myopia model to confirm the sensitivity and robustness of the protocol. Additionally, we show the OMR is sensitive to the contrast and temporal resolution of the sinusoidal gratings. Furthermore, we found that the time between stimulus presentations impact the spatial-frequency tuning function likely as time is required for zebrafish to return to baseline swimming after eliciting the OMR. Finally, we found that the OMR after ten versus twenty seconds of stimulus onset appears comparable; indicating that robust OMR responses in zebrafish can be elicited through relatively short stimulus presentations. Through the experiments conducted, we present an optimized protocol specific to zebrafish. The protocol may be used to follow the progression or treatment efficacy of progressive neurological disorders including specific visual disorders and higher brain functions with visual endophenotypes. Ultimately, this protocol allows for high-throughput robust measures of visual and neural function in zebrafish.

## Introduction

Vision is dependent on the integrated function of the eyes and visual pathway. Aberrant vision can be caused by abnormal ocular development (Chang et al., 2025; Janarthanan et al., 2025; Shandiz et al., 2011), injury or loss of retinal cells (De Silva et al., 2024; Grover et al., 1999), or neurological impairments (Bulens et al., 1986; Chen et al., 2005). As the retina is part of the central nervous system, neurological conditions can have visual endophenotypes (London et al., 2013; Marchesi et al., 2021), which can act as a biomarker to uncover or track underlying defects in brain circuit function (during disease progression or to assess treatment strategies).

Subtle visual deficits can be revealed in humans and animals using a range of non-invasive psychophysical tests. In the laboratory, visual responses can be elicited in animal models through methods that leverage visually evoked reflexes. Commonly used examples of approaches that tap into reflex visual pathways are the optokinetic reflex (OKR) (Cahill & Nathans, 2008; Zhu et al., 2025; Cameron et al., 2013; Gresty, 1975; Q. Shi & Stell, 2013; Tappeiner et al., 2012; Portugues & Engert, 2009) and optomotor response (OMR) (K. LeFauve et al., 2021; Naumann et al., 2016; Portugues & Engert, 2009; C. Shi et al., 2018; Srinivasan, 2011; Warren et al., 2001; Xie, et al., 2019; Xie, Bui, et al., 2023; Xie et al., 2024). These related processes involve eye reflexes (OKR) or head/body movement (OMR) due to motion detection in the surrounding visual environment. These responses help to keep the visual image stable on the retina and are crucial for preventing motion blur allowing for high-resolution vision. As these processes are highly conserved across all vertebrate models, they are robust visual tests that may be used for high-throughput genetic disease modelling or treatment screening.

Both the optokinetic reflex and optomotor response utilize similar neural networks across vertebrates (Kretschmer et al., 2017; Matsuda & Kubo, 2021; Naumann et al., 2016). Both responses are primarily driven by direction-selective retinal ganglion cells (Kretschmer et al., 2017; Matsuda & Kubo, 2021; Naumann et al., 2016). Visual information is processed by the pretectum (a cluster of neurons in the mid-brain that are responsible for integrating optic flow information). At this point, the neural circuity involved diverge to control either eye movements (OKR) or body motion (OMR). Eye movements to recover retinal position (recovery saccades) are driven by motor neurons which control the extraocular muscles (i.e., the lateral rectus and medial rectus) mediating sideways eye movements (Portugues et al., 2014). In contrast body movement in the OMR is driven by locomotive circuits which control head and body movement (Naumann et al., 2016).

OMR and OKR have been used for visual assessment in humans (Warren et al., 2001) and a range of vertebrate species, including guinea pigs (Jnawali et al., 2021), chicks (Diether & Schaeffel, 1999), mice (Kretschmer et al., 2017; C. Shi et al., 2018), insects (Srinivasan, 2011), and fish (K. LeFauve et al., 2021; Naumann et al., 2016; Portugues & Engert, 2009). One animal model that has been increasingly used for large-scale biomedical disease modelling (in part due to the ease of performing genetic manipulations) and treatment screening (high throughput swimming exposure) is the zebrafish (*Danio rerio).* Zebrafish have emerged as a key genetic vertebrate model organisms as they are particularly useful for their ease of care and breeding, large clutch sizes and rapid visual development (Crouzier et al., 2021; Renninger et al., 2011). Zebrafish share over 80% of disease-causing genes with humans, as well as conserved retinal and reflex pathway architecture. Hence, they are widely used in vision and neural research (Bilotta and Saszik, 2001; Gestri et al., 2012).

OMR may be used in preference to OKR to assay visual acuity in zebrafish as it requires less investigator intervention and carries lower animal welfare impacts (Kretschmer et al., 2017; C. Shi et al., 2018). OKR requires the immobilization of the animal of interest so eye motion can be more easily captured, while OMR allows for models to be unrestrained and their head/body movement to be tracked. Zebrafish utilize the optomotor response when viewing shadows along the riverbed to stabilize themselves within water (Holman et al., 2023). Hence, to elicit the response, scientists show larval and adult zebrafish moving visual stimuli (generally stripes) on a screen adjacent to (Karaduman et al., 2023; Maaswinkel & Li, 2003) or beneath the fish (Gore et al., 2024; Holman et al., 2023; K. LeFauve et al., 2021; Matsuda & Kubo, 2021; Naumann et al., 2016; Saputra et al., 2024; Xie et al., 2024). The zebrafish are said to have a positive response if they voluntarily swim in the direction of the grating movement. The majority of experiments have been performed using larval zebrafish (Holman et al., 2023; Maaswinkel & Li, 2003; Matsuda & Kubo, 2021; Naumann et al., 2016; Portugues & Engert, 2009; Xie et al., 2021). In contrast, fewer studies have considered OMR in adult zebrafish (Gore et al., 2024; K. LeFauve et al., 2021; Karaduman et al., 2023; Saputra et al., 2024) due to greater behavioural complexity reflected in swimming behaviour of adult fish. As such experimental set-ups that return robust measures of OMR in freely swimming adult fish have yet to be developed (Gore et al., 2024; K. LeFauve et al., 2021; Karaduman et al., 2023; Saputra et al., 2024). The capacity to quantify vision using OMR across the fish lifespan opens up the possibility to study chronic and progressive neurological and visual disorders, e.g. myopia (Baird et al., 2020) and glaucoma (Sheybani et al., 2020; Weinreb et al., 2014).

As zebrafish eyes are positioned laterally, stimuli positioned below is less optimal for adult fish. Here we describe a protocol, that optimises the OptoDrum (Striatech) to assay OMR in adult zebrafish. The device, originally designed for use with rodents, has four LED screens where rotating grating stimuli are displayed laterally around a central platform. In this paper, we present a protocol (animal setup and stimulus parameters) for assessment of OMR across a range of zebrafish ages (between 4 and 8 weeks of age). To do so, we measured the visual acuity of an environmental-myopia model at 6- and 8- weeks of age against wildtype zebrafish to assay protocol sensitivity and identify spatial frequencies which elicit positive optomotor responses. We optimised various parameters of the protocol by assessing performance with differing interstimulus rest periods (7 wpf), stimulus duration (4 wpf), contrast (9 wpf), and temporal frequencies (6 wpf). As such, we demonstrate how the OptoDrum could be used to measure the optomotor response in juvenile and adult zebrafish.

## Materials & Supplies

Zebrafish are maintained and bred at the *Danio rerio* University of Melbourne Fish Facility. Embryo and larval zebrafish are grown in petri dishes within a 28.5°C incubator up to 5-days post-fertilization (dpf). After which, zebrafish are transferred to tanks in a flow-through Tecniplast system. Fish are raised in a 14-hour/10-hour light/dark (L/D) cycle at 28°C. All procedures performed followed the code of practice for the care and use of animals by the Australian National Health and Medical Research Council. The study was approved by the University of Melbourne’s Large and Exotic Animal Ethics Committee (Project no. 30956).

Optomotor responses were compared at different contrasts, temporal frequencies, and interstimulus rest periods. All experiments were performed using wildtype AB zebrafish. Additionally, we used a previously established dark-rearing myopia (short-sightedness) model (Xie, Goodbourn, et al., 2023) to determine whether the protocol is robust enough to differentiate fish with different visual acuity. For the dark-rearing model, we transferred 4 wpf zebrafish to tanks wrapped in black cloth tape (Bear Brand, 66623336603) to block all light exposure for either 2- or 4-weeks. Twice daily health monitoring (∼ 1 min) occurred under dim red-light illumination (17.4 cd·m^−2^; λ_max_ = 600 nm). This group was compared to age-matched controls raised under standard L/D conditions.

## Detailed Methods

### Set-Up

#### Computer Set-Up

This study utilized the Striatech OptoDrum (Tübingen, Germany; L × W × H: 53 × 53 × 30 cm^3^) to assay OMR in zebrafish. The OptoDrum consists of four computer screens that surround a central platform (Figure 1A). During trials, vertical grating width is automatically adjusted to maintain the angular resolution from the central platform (e.g. widened towards the corners of each monitor). Two built-in programs, provided by Striatech, are primarily used to do this: OptoDrum.exe (version 17.1.1, Striatech; Figure 1B) and Demo Stimulus (version 2.0.1, Striatech; Figure 1C). Our study utilizes the Demo Stimulus Program to control grating parameters as it allows for hands-off, timed trials through customised JSON file.

**Figure 1.**
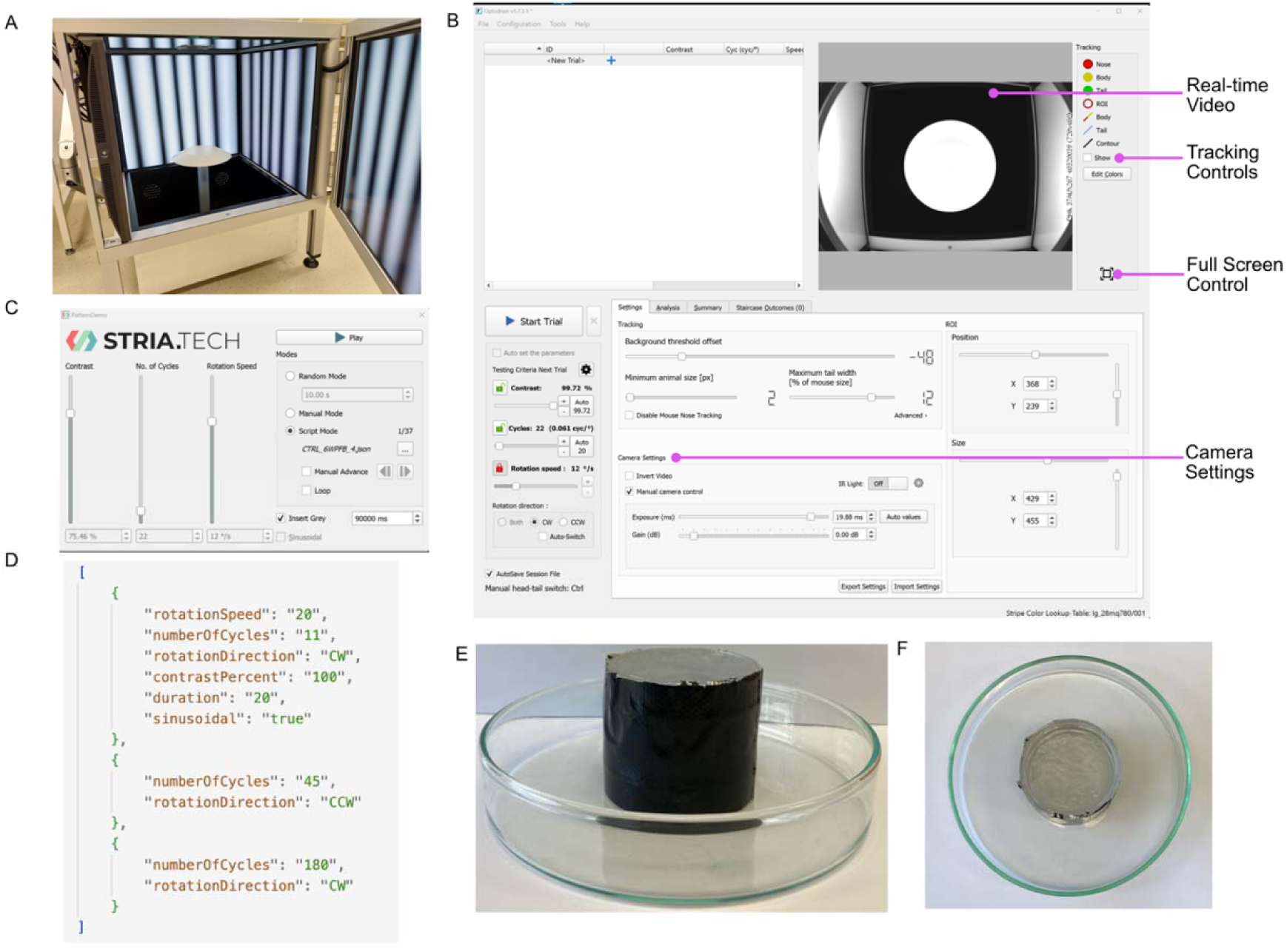
OptoDrum & zebrafish arena set-up. (A) Image of the OptoDrum with four LED computer screens displaying sinusoidal gratings around a central platform. (B) Image of OptoDrum.exe program that is native to the OptoDrum computer. Arrows indicate camera and video controls. (C) Demo Stimulus program used to display gratings to zebrafish. Script mode allows for selection of a json file. (D) Representative JSON file which would display three trials. Index 0 (row 2-9) controls the parameters for the first trial. It specifies the temporal frequency (rotation speed), spatial frequency (numberOfCycles), direction (rotationDirection; either clockwise [CW] or counterclockwise [CCW]), and trial duration in seconds. Gratings may be made sinusoidal (i.e. “sinusoidal: “true”). Index 1 (rows 10-13) and index 2 (rows 14-17) shows the parameters of trial 2 and 3, respectively. Only conditions that change between trials need to be specified in each index. Script must be enclosed in brackets (row 1and 18). (E,F) Images of the (E) side-view and (F) top-down view of the zebrafish arena crafted from a 14 cm glass petri and internal column of petri dish lids covered in black cloth tape.

During a trial, we record fish movements using the OptoDrum.exe program (up to 1 hour and 11 minutes). The program shows a live feed from a camera situated over the central OptoDrum platform, affording a view of the whole arena. The focus was adjusted to optimised clear viewing of the zebrafish. The Snipping Tool Windows Application (version 11, Microsoft) was used to screen-record the camera feed, for post-hoc analysis.

Stimuli were controlled via a JSON file (generated by a python program) to input into the script-mode of the Demo Stimulus program. For our experiments, the JSON file contained the temporal frequency (between 20 to 100 °/s), number of cycles (between 5 to 180 cycles), rotation direction (clockwise or counterclockwise), contrast (0 to 100%), and duration of each stimulus (20 seconds) (Figure 1D). The presentation order of the various stimuli combinations was randomised. Sinusoidal gratings were used for all experiments. A grey screen was displayed between trials for 30 to 90 seconds (dependent on experiment).

#### Swimming Chamber Preparation

The swimming chambers consists of a glass petri dish (SciLabware 1480/12D) with a 32 cm high central column. This inner central column consists of four 5.5 cm petri dish lids glued together by Aquarium Silicone Sealant (Sika, Catalogue No. 020515060000000003) and wrapped in cloth tape. The column was then glued to the centre of the 14 cm wide x 2.5 cm deep glass petri dish to create the swimming arena (Figure 1F). To remove any possibly toxic glue residue from the arena, we soaked the swimming chamber in water for 48 hours prior to use with zebrafish.

To prepare the arena for OMR, 150 mL of 28.5°C fish facility water was transferred to the circular arena (∼1.4 cm water height). This ensured sufficient height to comfortably cover the fish entirely, with the ventral fin clearing the bottom of the arena. For the OMR, individual fish are placed in the arena. A glass cover (15 cm glass petri dish) is placed over the arena to prevent any zebrafish from jumping out. The arena, with the zebrafish, was positioned at the centre of the OptoDrum platform directly under the recording camera. Throughout the experiment, the room is kept dark to prevent light reflections in arena water (which impedes fish tracking for downstream analysis). A five-minute acclimatisation period with an even grey (brightness of 144 lux) stimulus was displayed in the OptoDrum prior to visual stimuli presentations. During this time, fish were continuously monitored for any signs of distress or any other behavioural abnormalities (e.g. not moving, swimming erratically) that might exclude the individual fish from analysis. We did not encounter any such issues throughout any of our experiments.

Following conclusion of the trial, the fish were transferred back into 500 ml of fish water for subsequent humane killing 1000 ppm AQUIS (Ehrlich et al., 2019) and postmortem analysis (e.g. ocular coherence tomography, weighing, histological analysis of the retina and ocular structures using for example Haematoxylin and Eosin staining or immunohistochemistry, data not shown).

### General optimised OptoDrum protocol

Based on our results presented below, we propose following as a foundational protocol for optomotor response testing using the OptoDrum for zebrafish: After individual zebrafish are placed in the OptoDrum, they are allowed five minutes of acclimatisation time. The zebrafish are shown a random order of sinusoidal gratings of either 0.014 cycles/degree (c/°), 0.031 c/°, 0.061 c/°, 0.125 c/°, 0.250 c/°, or 0.500 c/° (which corresponds to 5, 11, 22, 45, 90, and 180 cycles), which spans the main spatial frequencies wildtype zebrafish respond to at these ages. Unless specifically assessing visual performance linked to contrast sensitivity or stimulus movement, the contrast (100%) and the temporal frequency (20 °/s) of the gratings can be kept constant for each trial. Between each trial a 90 second period of rest was used, during which a grey screen is shown. This allows fish to stop swimming in response to the previous stimulus. Usually, fish swimming returns to baseline swimming between 60 to 90 seconds. As such, 90 seconds is an adequate amount of time between stimulus presentations. For each stimulus the grating pattern can be shown for ten seconds as this is enough time for the zebrafish to observe the stimulus and elicit the OMR.

Throughout the entire trial, each spatial frequency is shown 3 times in the clockwise direction and 3 times in the counterclockwise direction. As such, each zebrafish is shown a unique set of 36 stimuli in a randomised order (see Supplementary File 1 for an example). Using this protocol each zebrafish trial runs for 1 hour and 5 minutes. Different comparisons were conducted to optimise these starting parameters as described in the Results section (Table 1). Of note, different aged fish were used to assess suitability of the protocol across ages.

**Table 1.**
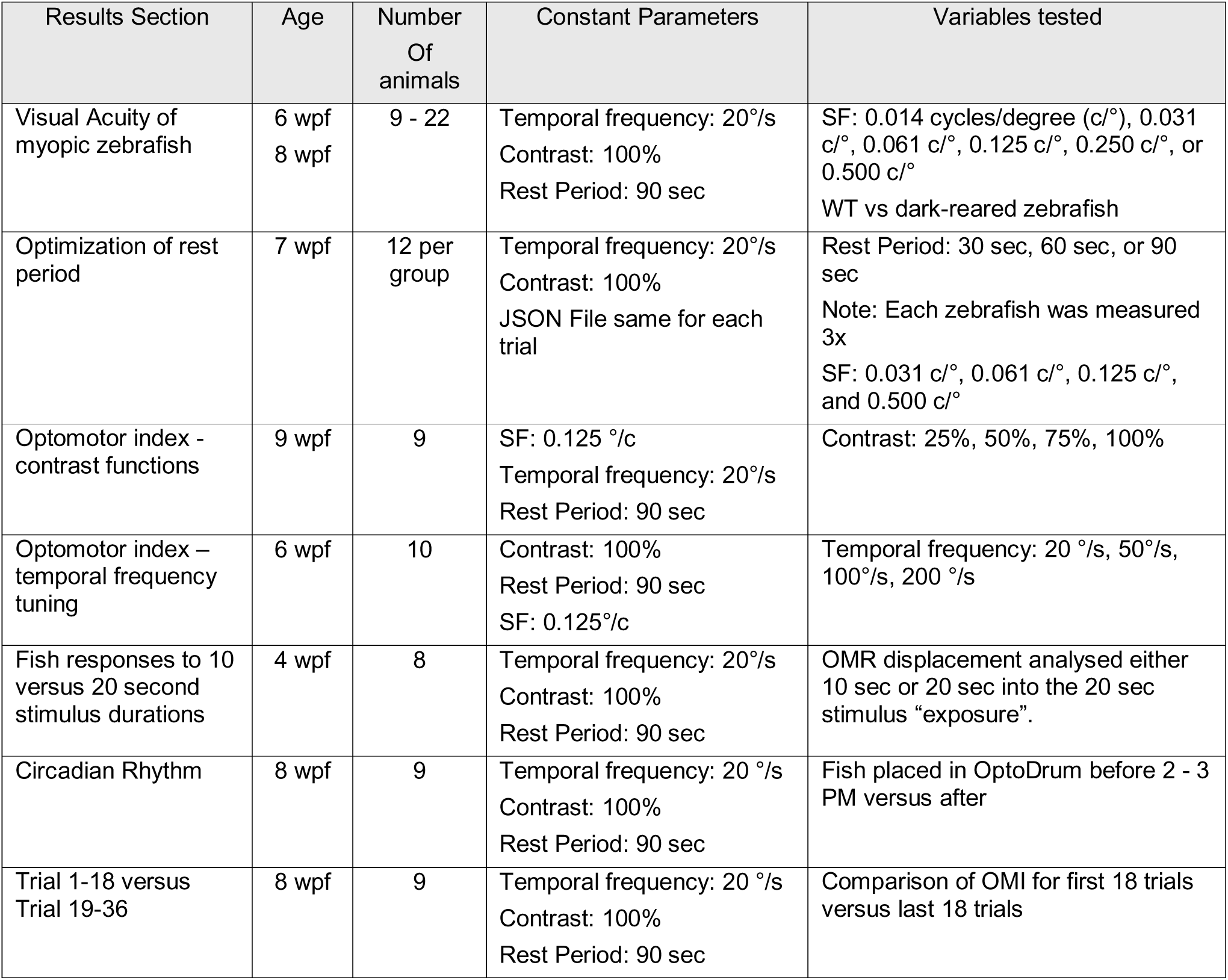
Summary of different parameter testing experiments. For most of these (unless stated otherwise) following parameters were kept constant: Spatial frequency = 0.014 cycles/degree (c/°), 0.031 c/°, 0.061 c/°, 0.125 c/°, 0.250 c/°, 0.500 c/°; contrast = 100%; interstimulus rest period = 90 seconds; temporal frequency 20°/s

### Data Analysis

Once the experiment has finished, the screen recording (captured by the snipping tool) was saved as a mp4 video file. Dependent on video length, it may be necessary to reduce the video file size for file transfer. To do so, we use the Handbrake application (version 1.9.2, Handbrake Team) to convert the mp4 video to a smaller video size.

The video file was separated into periods where the fish was shown stimuli from those periods without stimuli (i.e. grey screen). The stimulus clips are then cropped so only the fish swimming in the chamber can be seen on video — this aids fish tracking. Stimulus files are named based upon the parameters of the sinusoidal gratings shown.

All video processing was performed with the aid of custom python programs. Cropped stimulus videos were then processed by a published python package called Stytra (Štih et al., 2019). Like other tracking software, Stytra uses contrast to locate a zebrafish as it moves and outputs the x and y coordinates of the fish over time in a csv file. A custom MATLAB program utilizes the x and y coordinates of the fish to generate a trace of the fish movements and computes the optomotor index (OMI) for each trial. The OMI is the overall angular movement of the zebrafish in the direction of the rotating gratings. If the tracking software failed to track the fish (e.g. tracked a shadow or light reflection, fish too small for detection) then the OMI needed to be calculated manually. Of the 3,896 videos tracked in this paper, 777 were manually tracked. To do so, we would measure the angular momentum of the fish from the start position using FIJI (Schindelin et al., 2012). Any trial where the zebrafish did not move (i.e. OMI is equal to 0) was invalidated and excluded from analysis. We then normalized the results to the spatial frequency that illicit the largest response in the control fish to obtain the normalized OMI.

To create a spatial-frequency tuning function, the normalized OMI for the whole experimental group was plotted against spatial frequencies (MATLAB). For comparisons across groups, we fit a lognormal curve of the following equation:

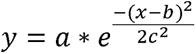

In the equation, y is the normalised OMI, a is the amplitude of the function, b is the peak spatial frequency, c is the standard deviation of the fit, and x is the log spatial frequency. The curve was fit using a least squares criterion using the Trust-Region algorithm.

### Statistical comparison

An omnibus F-test compares the goodness of fit (r^2^) between a full model (i.e. the parameter estimates between groups vary independently) to a restricted model (i.e. parameters are kept the same across groups). We then used a nested F_test_ to compare the full and restricted models (i.e. where one parameter is constrained across groups) to determine whether specific parameters (i.e. amplitude, peak spatial frequency, and bandwidth) differed significantly between groups (Lu & Dosher, 2013). A criterion of a = 0.05 was used to determine statistical significance.

## Results – Parameter optimisation

### Visual acuity of myopic zebrafish

To determine whether our OptoDrum protocol could discriminate OMR performance between groups of zebrafish with known visual deficits, we used our environmental-rearing myopia model (Xie et al., 2024). For this, 4 wpf zebrafish were dark reared for either two or four weeks (Figure 2A). After two and four weeks of treatment, we compared OMR performance of light/dark (L/D)-reared zebrafish versus dark-reared (D/D) zebrafish (6 wpf n = 22, 14, respectively; 8 wpf n = 9, 14, Figure 2). Two L/D-reared 6 wpf and one L/D reared 8 WPF zebrafish were excluded from analysis as they did not exhibit a measurable OMR (i.e. these zebrafish only swum in one direction for all trials). Experiments were performed over a two-day period once zebrafish were 6 and 8 weeks-post-fertilization using the generalized protocol, apart from a twenty second stimuli duration (as such, trial duration was 1 hour and 11 minutes).

**Figure 2.**
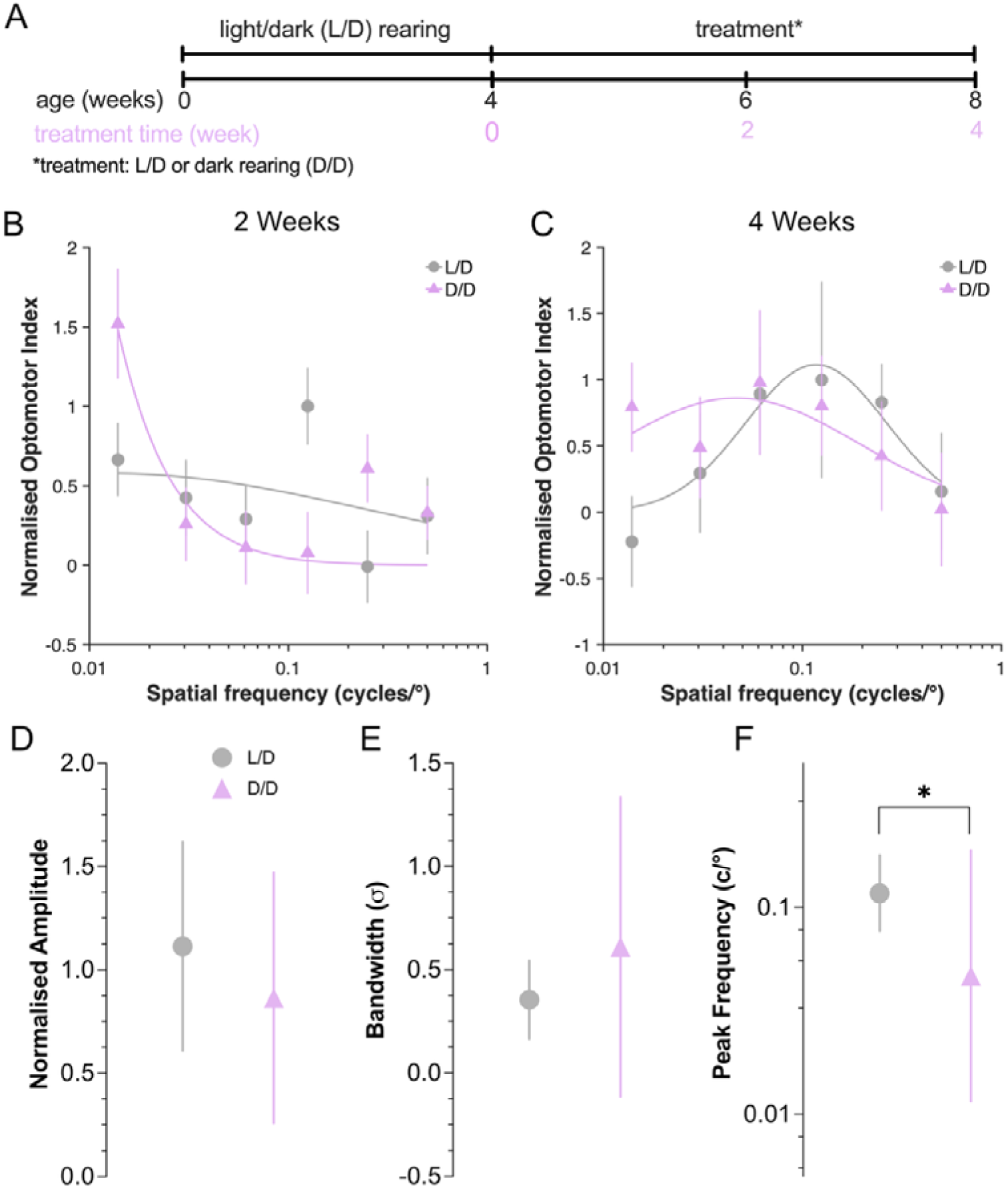
Spatial-frequency tuning functions of environmental-myopia zebrafish model. (A) Wildtype AB zebrafish were reared up to found weeks-post-fertilization in normal light/dark (L/D) conditions, consistent with standard husbandry. At which point, they were randomly assigned to either be dark reared (D/D) or kept under L/D conditions for either two or four weeks. (B, C) Spatial-frequency tuning graphs for fish reared under L/D or D/D conditions for (B) two (n = 22, n = 14, respectively) and (C) four (n = 9, n = 14, respectively) weeks. Sinusoidal gratings were shown at 20 °/s at 100% contrast. Curves are fit as three-parameter log-Gaussian functions to minimize the least-square error. (D-F) Comparisons of spatial-frequency tuning function parameters after four weeks of dark rearing for (D) normalised amplitude, (E) bandwidth, and (F) peak spatial frequency. Group data are shown as mean ± SEM in (B, C) and mean with 95% confidence intervals in (D–F). *p < 0.05 (F-test).

While at 6 wpf, we did not observe a log normal pattern commonly associated with spatial-frequency tuning graphs (Figure 2B), we did observe the expected pattern at 8 wpf (Figure 2C). We performed an omnibus fit test to compare the fit of the spatial-frequency tuning graphs for both timepoints and found no significant difference (Supplementary Table 1). We went on to assess whether there were significant differences in any parameters, including graph amplitude, bandwidth, and peak spatial frequency (6 wpf - 2 weeks treatment shown in Supplementary Table 1; 8 wpf - 4 weeks treatment shown in Figure 2D-F). While we found no significant difference in normalized amplitude or bandwidth, a significant shift occurred in peak spatial frequency at both treatment time-points (two weeks: F[1,6] = 21.63, p = 0.004; four weeks: F[1,6] = 6.27, p = 0.046). Only data from four weeks of treatment (8 wpf) is shown on Figure 2D-F; parameter estimates after two weeks treatment (6 wpf) is in Supplementary Table 2.

### Optimization of rest period

We tested whether the interstimulus rest period between trials might impact the spatial-frequency tuning curve. We hypothesised that longer rest periods between each stimulus trial would be beneficial for the zebrafish to return to baseline activity and remove the possible impetus to keep swimming in the same direction as per previous stimulus. To assess the minimum rest period duration required, experiments were performed over a three-day period when zebrafish (n = 12) were between 48 and 50 days-post-fertilization. Fish were randomly assigned to one of four groups, which would determine the order they were placed in the OptoDrum and sequence of 30-, 60-, or 90- second rest period between trials (Table 2). Once we had cycled through all four fish for Round 1, we placed Zebrafish 1 back into the OptoDrum for Round 2 etc. As such, each fish had been assayed three times in total over the course of one day with each of the rest period durations.

**Table 2.**
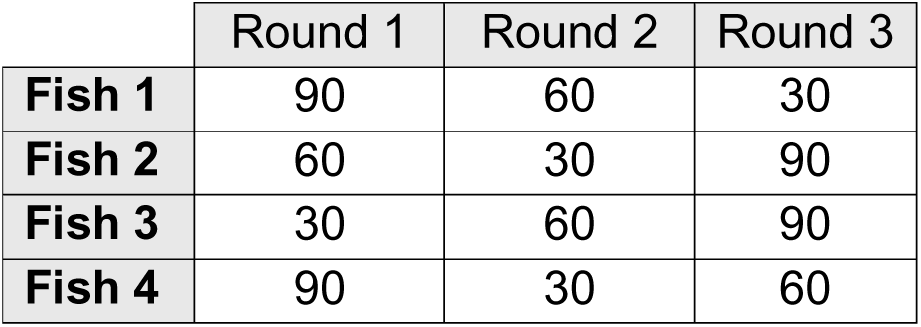
The order of zebrafish assayed to assess impact of between stimulus rest period duration on optomotor response. Numbers show duration of no-stimulus period in seconds.

|  | Round 1 | Round 2 | Round 3 |
| --- | --- | --- | --- |
| **Fish 1** | 90 | 60 | 30 |
| **Fish 2** | 60 | 30 | 90 |
| **Fish 3** | 30 | 60 | 90 |
| **Fish 4** | 90 | 30 | 60 |

The same JSON script was used for all fish and experimental rounds (Supplementary File 2). The stimuli displayed was randomly ordered using a custom python script. Sinusoidal gratings were only shown at the most responsive spatial frequencies of 0.031 c/°, 0.061 c/°, 0.125 c/°, and 0.500 c/° to reduce trial duration. Each of the 24 stimuli were presented for 20 seconds, resulting in trial durations of 25-, 37-, or 49-minutes for rest periods of 30, 60, or 90 seconds, respectively (including the initial five-minute acclimatisation time).

We found a significant difference between the spatial frequency tuning-curves of the 30- and 90- second rest curve from an omnibus fit test (F(3,2) = 54.33, p = 0.018; Figure 3, Supplementary Table 3). There was a significant difference in amplitude (F(1,2) = 94.15, p = 0.010), bandwidth (F(1,2) = 27.41, p = 0.035), and peak spatial frequency between the 30- and 90- second rest period curves. Of note, confidence intervals of the gaussian fit around the best fit parameters becomes large with shorter rest durations. This suggests that rest period of > 60 seconds provide better responses.

**Figure 3.**
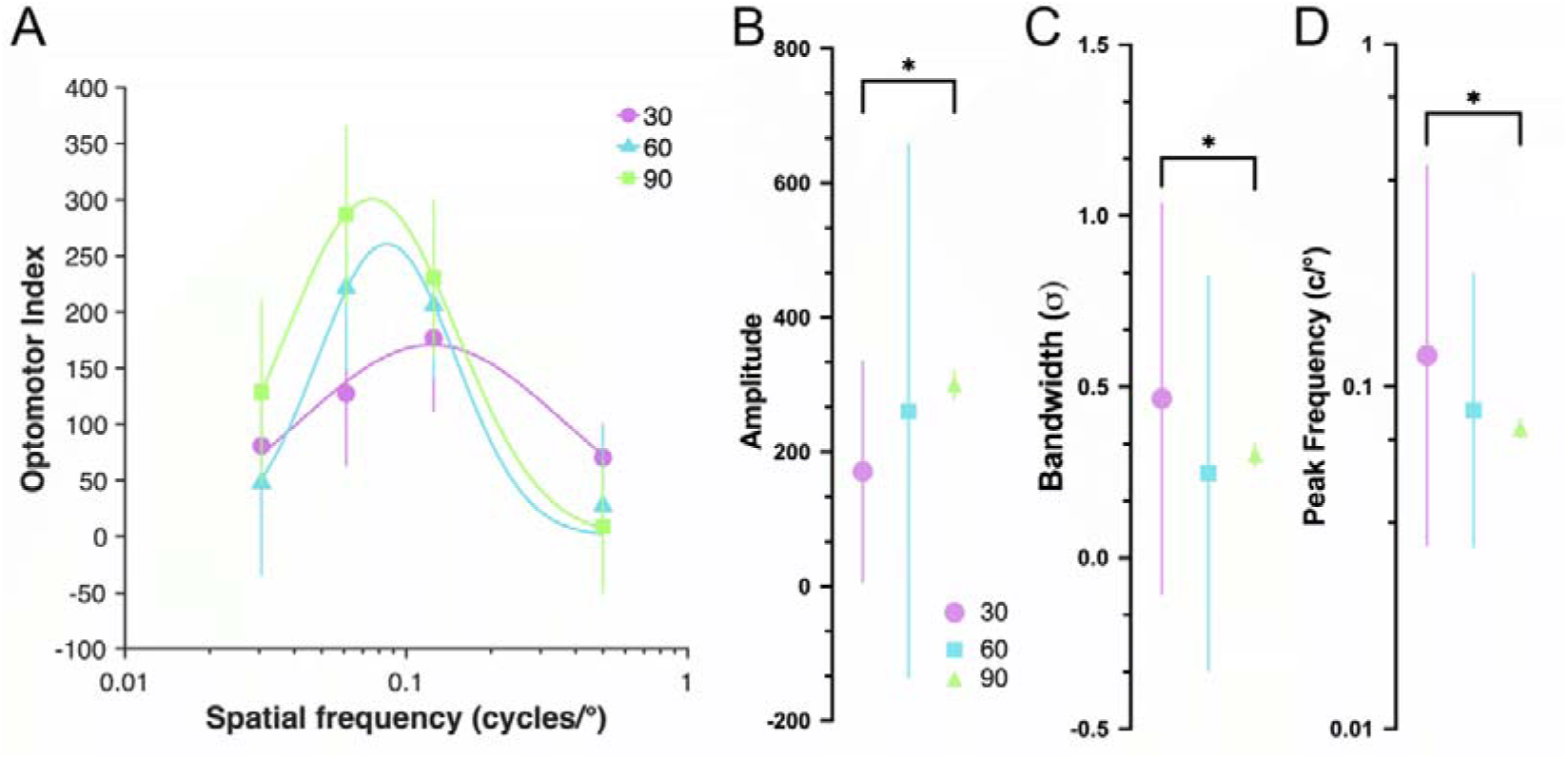
Spatial-frequency tuning functions of wildtype zebrafish with different rest period intervals between trials. (A) The spatial-frequency tuning function of 12 Wildtype AB zebrafish was found. Each zebrafish was placed in the OptoDrum on three separate occasions over a day period. The only parameter that changed between experimental rounds was the time interval between trials — either a 30-, 60-, or 90- second period of no-stimulus was used between trials. Sinusoidal gratings were shown at 20 °/s at 100% contrast. Curves are fit as three-parameter log-Gaussian functions to minimize the least-square error. (B-D) Comparisons of spatial-frequency tuning function parameters for (D) amplitude, (E) bandwidth, and (F) peak spatial frequency. Group data are shown as mean ± SEM in (B, C) and mean with 95% confidence intervals in (D–F). *p < 0.05 (F-test)

### Optomotor index - Contrast function

While most of our experiments are conducted at 100% contrast, various visual phenotypes might show changes in contrast sensitivity. Thus, we tested how changing contrasts might affect optomotor response behaviour. We measured 10 wildtype zebrafish aged at 9 wpf (66 dpf and 67 dpf). One of the zebrafish assayed was excluded from analysis as it only swum in one direction for all stimuli.

To assay contrast sensitivity, we generated JSON scripts which randomly allocated a contrast of 25%, 50%, 75%, or 100% contrast for each trial. The direction of stimuli was alternated, where the first stimuli was randomly chosen to be clockwise or counterclockwise. Stimuli were presented at 0.125 °/c (i.e. spatial frequency which illicit the largest OMR) for twenty seconds (90 second intervals in between stimuli). Trial duration was 49 minutes, including five-minutes of acclimatisation time.

As expected with higher contrast levels (75%, 100%) zebrafish exhibited more robust OMR compared with lower contrast stimuli (25%, 50%). This indicates that the OptoDrum may be used to assay changes in zebrafish contrast-sensitivity functions.

**Figure 4.**
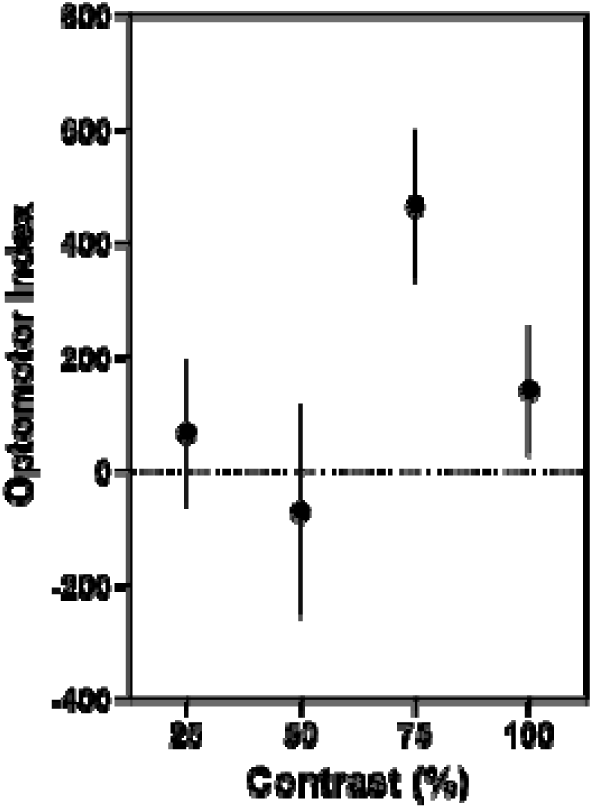
Optomotor index versus contrast for wildtype zebrafish. 9-week-old zebrafish (n = 9) were shown sinusoidal gratings at four different contrasts. Gratings had a spatial frequency of 0.125 cycles/° and temporal frequency of 20°/s. Group data is shown as mean ± SEM.

### Optomotor Index – Temporal frequency function

Temporal frequency of the stimuli may be used to test the temporal resolution of the visual system. Here, we tested how 6 wpf zebrafish respond to 20 °/s, 50 °/s, 100 °/s, or 200 °/s temporal frequencies for 0.125 c/° sinusoidal gratings. The direction of the trials was alternated between clockwise and counterclockwise trials, with the direction of the first trial being randomized. As such, 10 zebrafish were shown 6 repeats of each temporal frequency (3 in the clockwise direction and 3 in counterclockwise) for a total of 24 stimuli presentations (twenty seconds with 90 second rest period), resulting in a total trial duration of 49 minutes, including acclimatisation time.

We found that the 6-week-old zebrafish did not respond well to the 20 °/s temporal frequency gratings (contrary to our prior tests) but had strong OMRs at 50 °/s, 100 °/s, and 200 °/s (Figure 5). With the optimal temporal frequency appearing to be 100 °/s. However, it should be noted that the OptoDrum prevents combinations of certain high velocities and high spatial frequencies to prevent corkscrew patterns which does limit certain temporal frequencies from being tested. Thus, different parameter combinations might be most suitable to different experiments focusing on distinct visual parameters that might be linked for instance to neural endophenotypes being assessed in a particular study.

**Figure 5.**
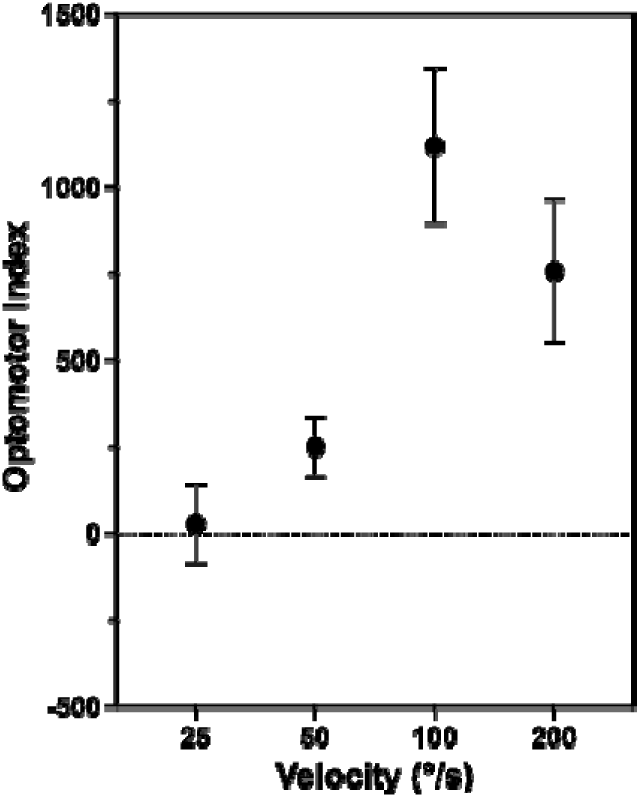
Optomotor index versus temporal frequency for wildtype zebrafish. 6-week-old zebrafish (n = 10) were shown sinusoidal gratings moving at four different velocities. Gratings had a spatial frequency of 0.125 cycles/° and 100% contrast. Group data is shown as mean ± SEM.

### Visual stimulus duration

Stimulus presentation duration may affect robustness of the optomotor response behaviour. On one hand a longer stimulus presentation allows fish to recognise the novel stimulus and compute whether to swim, and/or which direction, on the other hand fatigue may occur, where the initial drive to swim alongside the moving stimulus might be weaker after a while. We assessed this using our results from 4 wpf zebrafish (n = 12). Experiments were conducted over a two-day period using our generalized protocol, except for using a twenty second stimuli duration. All trials were manually tracked, where each video would be watched by two different individuals. The trackers would calculate the OMI at ten- and twenty-seconds and the calculated values would be averaged per video. Four zebrafish were excluded from analysis as they moved in one direction for all trials.

These analyses resulted in spatial-frequency tuning curves showing the OMI after 10- or 20- seconds of stimulus presentation for the same set of trials (Figure 6). We found that the spatial-frequency tuning graphs did not differ (F[3,6] = 0.04, p = 0.989, omnibus F-test). Comparisons of parameters found no difference in curve amplitude (F[1,6] = 0.10, p = 0.766, nested F-test), peak spatial frequency (F[1,6] = 0.00, p = 0.995), or bandwidth (F[1,6] = 0.01, p = 0.915). This indicates that trial duration may be reduced with limited impact to the normalized OMI. It suggests that the swimming behaviour of the fish is initiated relatively soon following stimulus onset. Additionally, once the zebrafish have picked a direction; they continue to swim in chosen direction for the duration of the trial. So, the primary difference between 10 and 20 seconds is the potential swimming distance, with no additional information gain.

**Figure 6.**
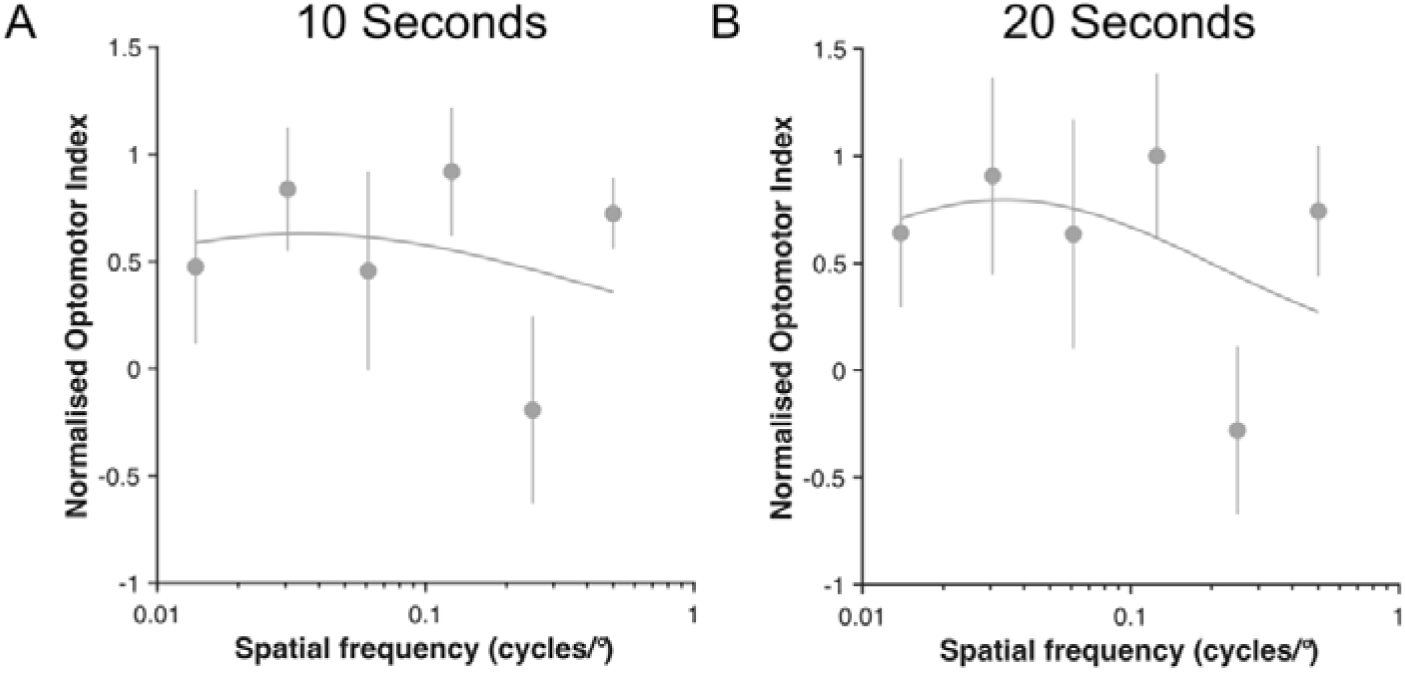
Spatial-frequency tuning graphs after 10- and 20- second trial duration. 4 WPF wildtype AB zebrafish (n = 8) were shown sinusoidal gratings of different spatial frequencies at maximal contrast (99.2%) and a 20 °/s temporal frequency. The optomotor index was calculated at (A) 10 seconds and (B) 20 seconds for the same set of trials. Curves are fit as three-parameter log-Gaussian functions to minimize the least-square error. Group data are shown as mean ± SEM.

### Time of Day Impact on OMI

As swimming behaviour can be influenced by the circadian rhythm (reviewed in Krylov et al., 2021), we compared the spatial-frequency tuning functions of fish run early versus late in an experimental day (Figure 7). To do so, we used data collected in the ‘Visual Acuity of Myopic Zebrafish’ section. Specifically, that of the wildtype zebrafish aged 8 WPF. We found the average time in which the zebrafish were placed into the OptoDrum was 2:30 PM. Any fish placed into the OptoDrum prior to 2:30 PM was labelled as ‘early’ (n = 5) and any after were considered ‘late’ (n = 4). No significant difference was found between spatial-frequency tuning curves from an omnibus F-test and no significant difference was found in gaussian parameters (amplitude, peak spatial frequency, bandwidth).

**Figure 7.**
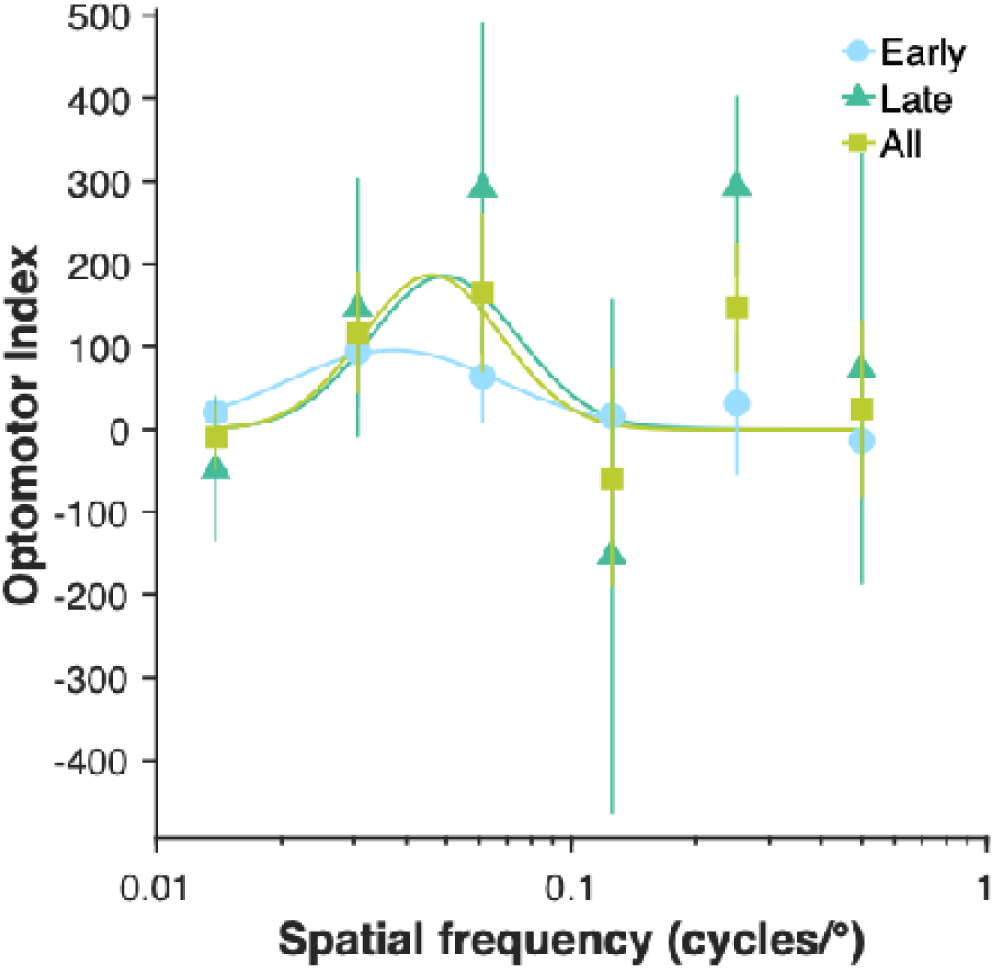
Optomotor index of zebrafish in the first half (prior to 2.30 pm) versus second half (after 2.30 pm) of an experimental day. 8 wpf wildtype AB zebrafish (n = 9), reared under standard lighting conditions, were shown sinusoidal gratings with 100% contrast and a 20 °/s temporal frequency. Zebrafish were classified as being early if experimental start time was prior to 2:30 PM (n = 5), and late if placed in OptoDrum after 2:30 PM (n = 4). (A) Spatial-frequency tuning curves of fish separated by time assayed in experimental day. Specifically, early in experimental day (n = 5), late in the experimental day (n = 4), and all fish measured during the experimental day (n = 9). Group data are shown as mean ± SEM.

## Potential Pitfalls & Trouble Shooting

### Set-up

While the OptoDrum screens show gratings at specific spatial frequencies, the change in refraction as light moves between one medium to another could distort the spatial frequency seen by the fish. Hence, while the protocol may be used to compare responses across different groups, it may not be able to deduce the exact limits of the zebrafish’s vision accurately. As the distance of the fisheye to the glass petri dish wall can also change, the fish’s location at any given time will further affect spatial frequency. To reduce variability associated with this, researchers could create different arenas for different age-groups. Lane size/width could be chosen as a function of zebrafish body mass/size. While narrower lanes restrict the range of spatial frequencies the fish may observe, lanes do need to be wide enough for good animal welfare. Zebrafish must be able to turn and move freely and not be stressed due to confinement.

For our experiments, we used a five-minute acclimatisation period. In this time, we monitor the fish for signs of distress and any behavioural abnormalities. Occasionally, zebrafish would not move during the acclimatisation time and only exhibit the OMR after a few trials. It may be beneficial to use *test* stimuli (i.e. show gratings that elicit a robust OMR) to confirm zebrafish behaviour before proceeding to experimental trials.

Individual zebrafish can display periods of only clockwise or counterclockwise swimming, which may be linked simply to swimming momentum. In our paradigm, we initially randomised (rather than alternated) direction of stimuli when assaying visual acuity. To minimise this potential, we used a 90 second rest period between stimuli, providing the opportunity for the fish to return to basal movement prior to the next trial. Another option may be to pause trials until the fish has returned to baseline behaviour or is stationary. For the contrast function and temporal frequency function, we did use alternate direction of stimulus (rather than random). Alternating swimming stimuli may work, as it requires the fish to turn around every time (i.e. the same “effort” required to follow the stimulus vs. the potentially easier behaviour to continue swimming momentum in the same direction).

Additionally, zebrafish swimming behaviour may also be impacted by water temperature. Ideally, the water should be kept at approximately 28.5 °C. It is important to note that the OptoDrum monitors generate heat which could impact water temperature. When 23°C water was placed in the OptoDrum, it rose to 27.5°C over a five-hour period (Supplementary Figure 1).

### Stimulus parameters

In this paper, we measure the zebrafish optomotor response between 4 and 9 wpf. While parameters used in this experiment may work for these age groups, it is important to define specific ranges for spatial frequency, temporal frequency, and contrast with your age-group of interest. Older zebrafish have previously been shown to favour higher spatial frequencies compared to larval and juvenile zebrafish (K. LeFauve et al., 2021; Karaduman et al., 2023). While we wanted to compare the spatial resolution limits of zebrafish vision between treatment groups, this may be unnecessary for other experimental designs. Using more spatial frequencies would improve gaussian curve modelling at the expense of longer experimental time.

Greater variability in behaviour can arise due to difference in activity from circadian rhythms (Krylov et al., 2021). For young larval zebrafish, rapid developmental and neural circuit maturation could mean that optomotor performance may vary substantially even within an experimental day (Xie et al., 2024). Such developmental refinement of neural circuit function are unlikely contributors to performance at the later ages we are assessing here. Nonetheless, it may be important to restrict the experimental window to specific times in the day (Figure 7), and for group comparison to interleave fish from various treatment groups.

Additionally, it has been suggested that zebrafish will stop showing an OMR after a certain number of stimuli presentations (K. LeFauve et al., 2021). We found a general decrease in OMR in the second half a trial compared to the first half (Supplementary Figure 2). Whether this is a behavioural response (no benefit of performing OMR) or related to swimming fatigue needs to be determined. However, it does imply that using *a priori* knowledge and (hypothetical) predictions of which aspect of the visual stimulus is likely most relevant will allow researchers to minimise other parameters and minimise OMR duration, thus possibly preventing decline in responses with time, and also allowing for more animals to be tested on the same day.

### Tracking

For the purposes of our experiment, we did not require Stytra to perfectly track the fish. During analysis of our experiments, we did observe that re-analysis and manual tracking especially for the young larvae was required. Similarly, if more precise analysis is required, e.g. in cases where tail behaviour or similar types of behavioural movement are important, video production and quality could be improved. Optimising these aspects may also benefit those who want to track multiple fish at the same time, however note, that shoaling behaviour emerges by after 2 wpf (reviewed in Facciol et al., 2020).

For optimisation of the videos, exposure or video gain could be adjusted to increase the contrast between the fish and background. Various modifications from the rodent optimised setup in the OptoDrum can be employed, including for example moving the OptoDrum central platform towards the camera seems to allow for a better-quality video by adjusting fish size on the camera feed and or using black cylindrical shields (Supplementary Figure 3) to reduce artefact due to excess light from the monitor being reflected by the glass petri dishes. These changes should not impact the fishes’ view of the rotating gratings.

High quality mp4 videos can be generated using the Window’s snipping tool as the recording software. We used the handbrake application to convert these videos to a lower quality, which fitted our requirements and optimised data storage constraints. To prioritize quality, it is possible to separate trials from the rest periods on the OptoDrum computer using the original mp4 video. If processing must be performed on a different computer, the handbrake application allows for adjustments to the compression level of videos and the speed of its encoding which may benefit video quality (although would require more storage space and additional conversion time).

## Concluding remarks

The zebrafish has emerged as a powerful biomedical vertebrate model system including for studies of neural development (including ecotoxicological impacts), neural disease modelling and treatment screening, in part due to the ability to rapidly combine genetic manipulations and environmental exposures to study their interactions in vivo. Such studies include those focused on the visual system, but also those assessing complex multifactorial central nervous system traits that often display specific visual endophenotypes. As a non-invasive behavioural output, both optokinetic and optomoter responses are powerful screening methods. Whilst established well in very young larvae (< 7 dpf), robust methods are not yet well established for older fish, which may be required for longitudinal studies of phenotype progression, and comparison of novel therapeutic treatment interventions across time. Here we present optimised techniques as a reference for juvenile zebrafish optomotor behaviour using the OptoDrum. Specific parameters we tested here can be incorporated in similar OMR setups (including user made equipment) and will allow the field to use standardised protocols to support cross-study comparisons.

## Supplementary Information

### Supplementary Tables

**Supplementary Table 1.** Statistics of spatial-frequency tuning functions for dark-reared myopic zebrafish after two- and four- weeks of treatment.

|  | 2 weeks treatment | 4 weeks treatment |
| --- | --- | --- |
| Omnibus comparison | $F(3,6) = 1.08$<br>$p = 0.427$ | $F(3,6) = 2.67$<br>$p = 0.141$ |
| Amplitude comparison | $F(1,6) = 17.56$<br>$p = 0.006$ | $F(1,6) = 2.86$<br>$p = 0.141$ |
| Peak frequency comparison | $F(1,6) = 21.63$<br>$p = 0.004$ | $F(1,6) = 6.27$<br>$p = 0.046$ |
| Bandwidth comparison | $F(1,6) = 1.80$<br>$p = 0.229$ | $F(1,6) = 2.95$<br>$p = 0.137$ |

**Supplementary Table 2.** Parameter values of spatial-frequency tuning functions for dark-reared myopic zebrafish after two-weeks of treatment. Numbers are displayed as mean with the 95% confidence interval in brackets.

|  | L/D- Reared | D/D- Reared |
| --- | --- | --- |
| Amplitude | 0.5794 [-0.7928, 1.952] | 1.396e+04 [-1.808e+07, 1.811e+07] |
| Peak frequency | 0.01282 [ 9.931e-16, 1.655e+11] | 1.763e-07 [ 0, Inf] |
| Bandwidth | 1.284 [-10.02, 12.59] | 1.145 [-73.28, 75.57] |

**Supplementary Table 3.** Statistics of spatial-frequency tuning functions for wildtype zebrafish with different interstimulus rest periods.

|  | 30 vs 60 seconds | 60 vs 90 seconds | 30 vs 90 seconds |
| --- | --- | --- | --- |
| Omnibus comparison | $F(3,2) = 4.61$<br>$p = 0.184$ | $FF(3,2) = 5.69$<br>$p = 0.153$ | $F(1,2) = 1.23$<br>$p = 0.382$ |
| Amplitude comparison | $F(1,2) = 8.60$<br>$p = 0.099$ | $F(3,2) = 5.69$<br>$p = 0.153$ | $F(1,2) = 94.15$<br>$p = 0.010$ |
| Peak frequency comparison | $F(1,2) = 5.16$<br>$p = 0.151$ | $F(1,2) = 2.71$<br>$p = 0.242$ | $F(1,2) = 40.92$<br>$p = 0.024$ |
| Bandwidth comparison | $F(1,2) = 7.92$<br>$p = 0.107$ | $F(1,2) = 1.23$<br>$p = 0.382$ | $F(1,2) = 27.41$<br>$p = 0.035$ |

### Supplementary Figures

**Supplementary Figure 1.**
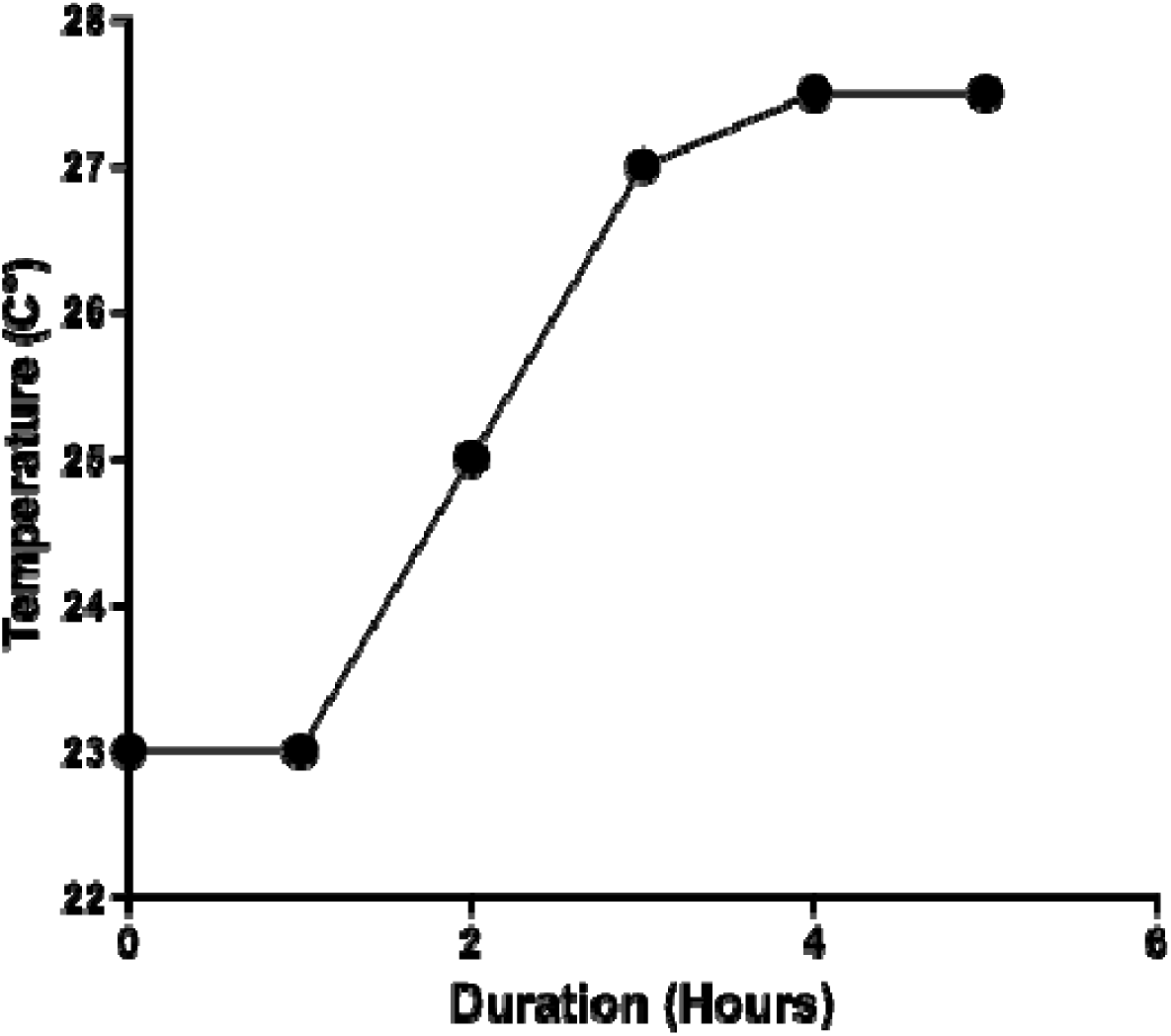
Temperature of water left in OptoDrum over 5 hours. 23°C water was placed in the OptoDrum while monitors were running. Every hour, the temperature of the water was measured for a total of five hours.

**Supplementary Figure 2.**
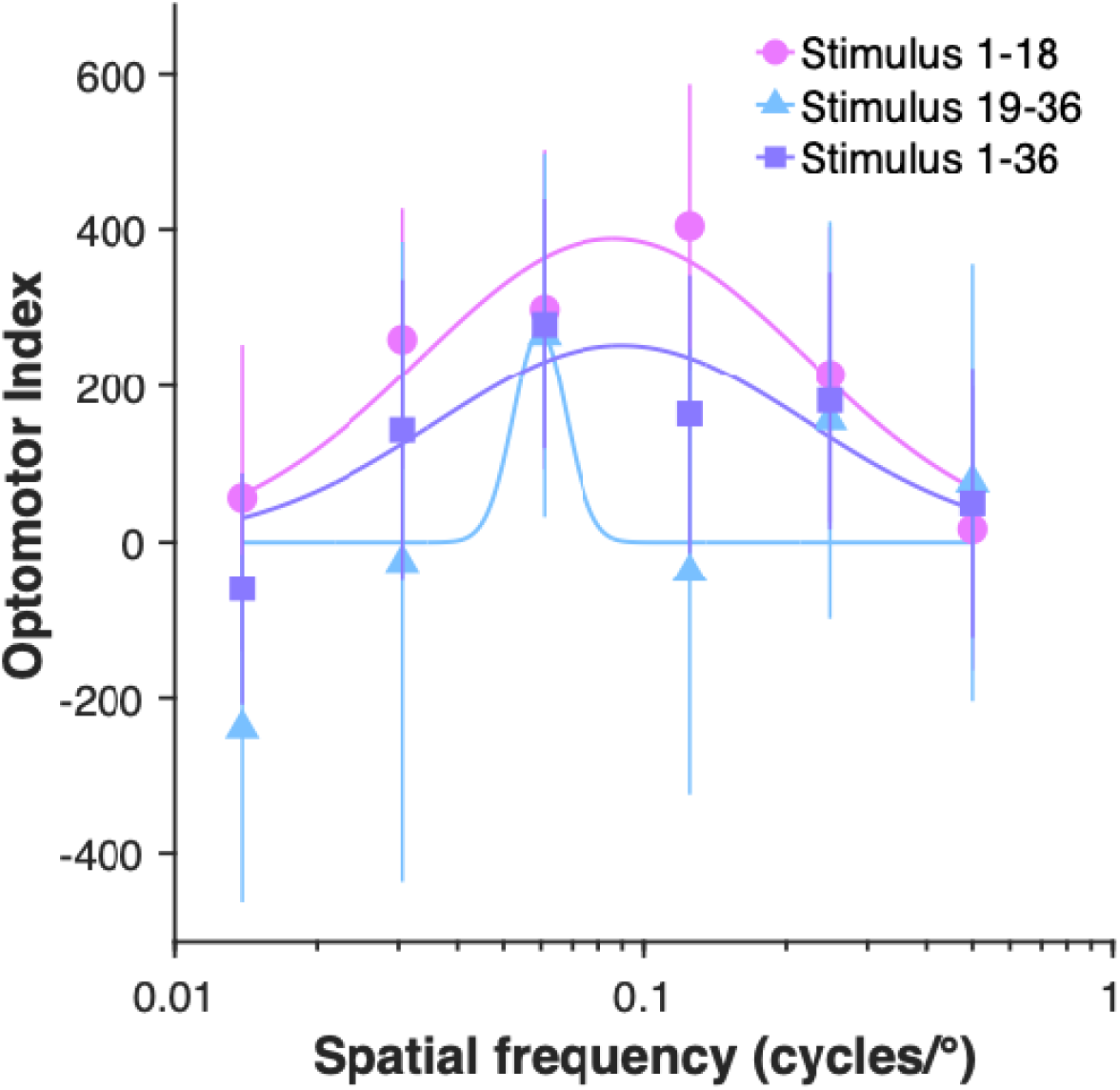
Spatial-frequency tuning function of zebrafish in the first half versus the second half of trials. 8-week-old wildtype AB zebrafish (n = 9) reared under standard lighting conditions were shown sinusoidal gratings with 100% contrast and a 20 °/s temporal frequency. Stimuli were separated based upon whether they were presented in the first half (1-18) or the second half of a fish trial (19-36). Fish were shown gratings at 0.014 c/° (n = 34 trials, 20 trials, respectively), 0.031 c/° (n = 31 trials, 23 trials, respectively), 0.061 c/° (n = 22 trials, 32 trials, respectively), 0.125 c/° (n = 23 trials, 31 trials, respectively), 0.250 c/° (n = 25 trials, 29 trials, respectively), and 0.500 c/° (n = 25 trials, 27 trials, respectively). The Stimuli 1-36 group show all trials (n = 54 per spatial frequency). Group data are shown as mean ± SEM.

**Supplementary Figure 3.**
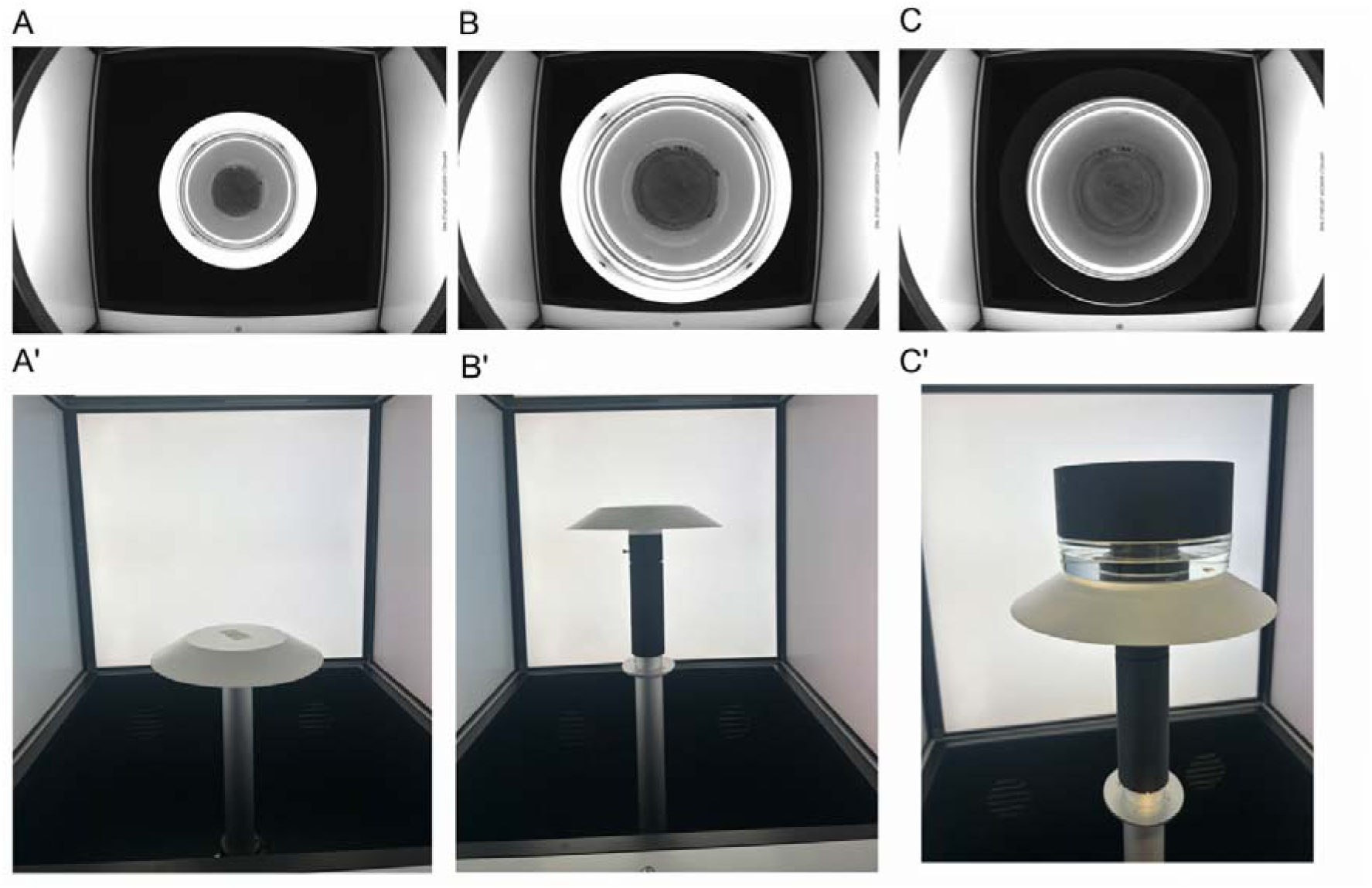
OptoDrum set-up options to optimize vision of zebrafish. Different arena designs were trialled to assess visibility of fish for automated tracking software. (A-C) Top-down images of a zebrafish in a glass arena (A) without additional platform elevation and a glass lid, (B) with platform elevation and a glass lid, and (C) with platform elevation and a cylindrical overlay placed atop arena. (A’-C’) Side view images of OptoDrum (A’) with normal set-up and (B, C’) with additional elevated platform. (C’) Zebrafish placed in arena with black cylindrical overlay placed atop arena.

### Supplementary Files

Note for submission the file extension was changed to “.txt”, for running on OptoDrum, simple change the file extension to “.json”

**Supplementary File 1.** Exemplar JSON file used for a fish measured in ‘visual acuity of myopic fish’.

**Supplementary File 2.** JSON file used when assaying differences in spatial-frequency tuning curves with different timed intervals between stimuli presentations.

